# Genomic diversity, population structure and admixture in native cattle breeds of Benin

**DOI:** 10.1101/2025.09.25.678660

**Authors:** Loukaiya Zorobouragui, Stephane R. Tapsoba, Amadou Traore, Kathiravan Periasamy, Rudolf Pichler, Tafara K. Mavunga, Alassan S. Assani, Hilaire S.S. Worogo, Nassirou Taba, Maximilien Azalou, Christophe Iwaka, Ibrahim T. Alkoiret, Isidore Houaga

## Abstract

This study investigates the genetic diversity, population structure, and admixture patterns of indigenous cattle breeds in Benin, shedding light on their evolutionary relationships and adaptation to the West African environment. A total of 348 cattle from eight indigenous breeds of Benin, including taurine (Lagune, Borgou, Pabli, and Somba), zebu (Gudali, Zebu Peuhl, and Yakana) and one crossbred (Bourgou X Zebu) cattle were genotyped along with a reference dataset of cattle from Europe, Asia, and West Africa. After quality control, 28.591 SNPs from 838 cattle were analyzed for genetic diversity, differentiation, and admixture. Pairwise F_ST_ values revealed significant genetic differentiation between local taurine and zebu breeds (F_ST_ = 0.05 – 0.15), with some populations showing close genetic relationships, while others, such as the Borgou and N’Dama breeds, exhibiting relatively more divergence. The admixture analysis indicated significant gene flow from zebu cattle into local taurine breeds, suggesting adaptive introgression driven by factors such as heat tolerance and disease resistance. Additionally, the effective population size (*N_e_*) was relatively higher in Benin’s taurine breeds as compared to zebu, likely attributable to traditional open communal mating practices. The genetic structure also reflected the influence of both historical and ongoing introgression from Asian zebu cattle. The results highlight the importance of maintaining genetic diversity through regional breeding strategies that consider environmental and adaptive pressures. The results of the present study will serve as a basis for the development of Community Based Breeding Programs (CBBPs) for Beninese cattle adapted to local contexts, integrating information on admixture levels, breeders’ preferences, production performance, and the conservation of local genetic diversity.

## Introduction

Cattle have historically played a pivotal role in global agriculture, serving as a primary source of milk[1], meat [2] and hide [3]. Beyond their economic value, they hold profound cultural and social significance, being integral to rituals and traditions associated with birth, marriage, death, and other ceremonial practices across several communities across the world. The domestication of cattle is traced back to two primary centers of origin: Eurasian taurine (*Bos taurus taurus*), domesticated in Eastern Europe, and Asian indicine (*Bos taurus indicus* or zebu), domesticated in the Indian subcontinent [4]. These domestication events, which occurred around 8,000 years ago, shaped the genetic variability observed in cattle today. Human migration and ecological adaptation facilitated the spread of cattle, subjecting them to various selection pressures that led to the development of distinct breeds with specialized traits for dairy, draft and meat production, and resilience to environmental stressors[5], [6], [7]. Over time, the evolutionary, ecological and management factors shaped the complex genetic architecture of cattle, resulting in extensive phenotypic and genetic diversity observed across global populations [8], [9], [10].

In Africa, the introduction and diversification of cattle followed distinct patterns, particularly in West and East Africa[11], where both taurine and zebu cattle have played significant roles[12], [13]. West Africa is home to a large number of taurine cattle, such as the Muturu, N’dama, Lobi and Lagune breeds, while in East Africa, taurine populations like the Sheko are now predominantly crossbred[14], [15], [16]. The arrival of first zebu cattle in Africa is believed to have occurred around 3,500 YBP, with subsequent admixture events significantly influencing the genetic makeup of African cattle populations[17], [18], [19]. In Benin, several cattle populations, estimated to be admixtures of taurine and zebu breeds, were introduced around 1,400 YBP [18], [20]. This background sets the foundation for a comprehensive study of Benin’s cattle breeds, in order to understand their genetic structure and evolution. The native cattle breeds of Benin comprise taurine and indicine breeds as well as their crosses. The taurine breeds namely Borgou, Somba, Pabli and Lagune, indicine breeds such as Gudali, Yakana and Zebu Peuhl and a taurine-indicine crossbred (Borgou X Zebu), provide an important genetic resource for investigating genomic diversity, population structure, and admixture. These breeds, integral to the livelihoods of smallholder farmers due to their resilience to transhumance pastoralism, diseases and harsh climates, are under increasing threat from indiscriminate crossbreeding with zebu cattle in the attempts to improve productivity[21], [22], [23], [24]. This crossbreeding threatens the genetic integrity of the native breeds and, combined with the absence of structured breeding programs and performance recording systems, exacerbates the vulnerability of these cattle populations. While some studies have explored crossbreeding and transboundary transhumance in African cattle[25], [26], [27], there is a significant gap in our understanding of the genomic diversity and population dynamics of Benin’s native breeds.

Genetic studies on African cattle populations have consistently revealed extensive admixture between *Bos taurus taurus* and *Bos taurus indicus*. For instance, Weerasinghe et al.[16] demonstrated the hybrid nature of cattle populations across East Africa using autosomal SNP markers. However, little research has been conducted on the genetic makeup of Benin’s cattle. The study by Vanvanhossou et al.[28] highlighted important morphometric diversity in Benin’s cattle. Vanvanhossou et al.[29] research on genetic admixture, inbreeding and selection signatures in five taurine breeds highlights the importance of understanding genetic diversity to ensure the sustainability of livestock production in Benin. Studies suggested that zebus were introduced to Benin about 1,400 YBP via the Indian Ocean trade[30], [31], [18], leading to the admixture observed today. However, there is limited knowledge on how this admixture has shaped the genetic structure and adaptive traits of Benin’s native cattle breeds. This study seeks to fill this knowledge gap by providing a comprehensive analysis of the genomic diversity, population structure, and admixture in native cattle breeds of Benin.

## Materials and Methods

### Animal Ethics statement

No animals were specifically sampled for the present study. All genotypic data analyzed were derived from existing DNA/sample repositories at Laboratory of Ecology, Health and Animal Production (LESPA), University of Parakou, Parakou, Benin. As no live animals were handled and no new samples were collected, ethical approval was not required for this study.

### SNP Genotyping

A total of 348 cattle belonging to eight indigenous cattle breeds including four taurine (Borgou, Lagune, Pabli, and Somba), four Zebu (Gudali, Zebu Peuhl, Yakana), and one crossbred (Borgou × Zebu) cattle from Benin were genotyped using Axiom Bovine Genotyping BovMDv3 array on the Affymetrix-Axiom GeneTitan Platform. In addition, genotype data of 100 Burkinabe cattle, 45 Malian cattle, 47 Nigerien cattle, 235 European taurine and 110 South Asian Zebu cattle, already available at Animal Production and Health Laboratory, Joint FAO/IAEA Centre, International Atomic Energy, Vienna, Austria from the same genotyping platform was utilized for comparative analysis. A summary of the dataset used in the study is presented in **Table 1**.

**Table 1:** Information on the cattle breeds studied and reference breeds.

| Country_Name | Species | Breeds names | Breeds_Code | Effectif |
| --- | --- | --- | --- | --- |
| Austria | European Taurine | Simmental | FLV | 60 |
| India |  | Jersey | IJR | 30 |
| Serbia |  | Holstein | SHO | 83 |
| Peru |  | Brown Swiss | YBS | 62 |
| Mali | West African Taurine | N'dama | AND | 45 |
| Benin |  | Borgou | EBG | 51 |
|  |  | Lagune | ELN | 29 |
|  |  | Pabli | EPB | 37 |
|  |  | Somba | ESO | 47 |
| Benin | Zebu X Taurine crossbred | Borgou X Zebu Crossbred | EBZ | 24 |
| Benin | West African Zebu | Beninese Gudali | EGO | 48 |
|  |  | Yakana | EYK | 31 |
|  |  | Beninese Zebu Peuhl | EZP | 81 |
| Burkina Faso |  | Burkinabe Bororo | FBO | 14 |
|  |  | Burkinabe Gudali | FGO | 45 |
|  |  | Burkinabe Zebu Peuhl | FZP | 41 |
| Niger |  | Bororo Niger Diffa | NBD | 21 |
|  |  | Bororo Niger Maoua | NBM | 26 |
| India | Asian Zebu | Ongole/Nellore | ION | 39 |
| Pakistan |  | Red Sindhi | PRS | 37 |
|  |  | Tharparkar | PTP | 34 |
|  |  | Total |  | 885 |

### SNP genotype data quality control

A total of 43,566 SNPs from 885 animals were received from the genotyping platform. The quality control was conducted using PLINK version 1.9 [32]. The following filtering criteria: (i) including animals that had a call rate higher than 0.95(ii) excluding SNPs corresponding to unknown chromosomes or sex chromosomes (iii) excluding SNPs that were genotyped in less than 90% of the samples. (iv) excluding SNPs with a minor allele frequency (MAF) less than 0.05 (5%), (v) excluding SNPs that deviated from Hardy-Weinberg equilibrium (--hardy: 0.000001 or 1e-6), (vi) excluding SNPs exhibiting high linkage disequilibrium (LD) (--indep-pairwise 50 5 0.5”) were implemented using PLINK 1.9 [32]. To identify potential monozygotic twins and duplicate samples, relatedness was evaluated using (--king: ∼ 0.354). After all the quality control, we were left with 28,591 SNPs and 838 cattle from 21 breeds for further downstream analyses.

### Data Analysis

#### Genetic diversity and population differentiation

Genetic diversity indices such as observed heterozygosity (Ho), expected heterozygosity (He), and the inbreeding coefficient (F_IS_) values were calculated using PLINK v1.9[32]. Principal component analysis (PCA) was performed to analyze ordinal relationships between populations and individuals using PLINK’s “--pca” function. In order to capture the genetic differentiation between cattle populations, we used the Hierfstat package as implemented in R [33]. The genetic differentiation of the breeds was examined by comparing their pairwise F_ST_ values.

#### Effective population size (Ne)

SNeP v1.1 based on linkage disequilibrium (LD), was implemented to estimate the trends in effective population size (Ne) of the studied populations [34]. The Ne estimations over different generations in the Beninese cattle breeds were visualized using a custom R script.

#### Admixture and ancestry analyses

The admixture analysis was performed for K values ranging from 2 to 10 [35] as implemented in the ADMIXTURE software [36] and visualized using R version 4.4.0 [37]. To determine the optimal number of ancestral populations supported by the data, the cross-validation error (CVE) approach was applied. This allowed the determination of the most suitable K-value for different likelihood runs using R. The K value with the lowest CVE was considered as the best fit [35].

#### Population splits tree and gene flow

A Neighbour-Net graph using pairwise Reynolds’ distances was computed using SplitsTree v4.14.6 [38]. To investigate potential gene flow patterns among the studied cattle breeds, TreeMix v1.13 [39] was utilized to examine the gene flow. Migration events were set from 0 to 5. A maximum likelihood tree representing the 17 African and Asian cattle populations, including admixture events, was constructed. Initially, a phylogenetic tree of these populations was built, and subsequently, migration events (modeled as edges) were incrementally added to the phylogenetic model. The migration edges were added until 98.68% of the variance in ancestry between populations was explained by the model. The residuals from fitted model were visualized using the R script implemented in TreeMix package.

#### Data availability

All relevant data are included in the manuscript. Additional data supporting the findings of this study, including the SNP genotype datasets generated, are available in the Supplementary Information (S1 file).

## Results

### Genetic diversity

Analysis of genomic diversity in indigenous cattle populations of Benin and comparison with other West African, European and South Asian cattle, revealed significant variability for observed heterozygosity (Ho), expected heterozygosity (He) and inbreeding coefficient (F_IS_). Estimates of genetic diversity indices (H_o_, H_e_ and F_IS_) are presented in **Table 2**. In African cattle breeds, H_o_ ranged from 0.259 to 0.356 with the lowest value recorded in the Lagune (ELN) taurine cattle of Benin and the highest value in the Bororo (FBO) zebu cattle from Burkina Faso. The expected heterozygosity ranged from 0.269 (ELN, Benin) to 0.339 (EZP, Benin).

**Table 2:**
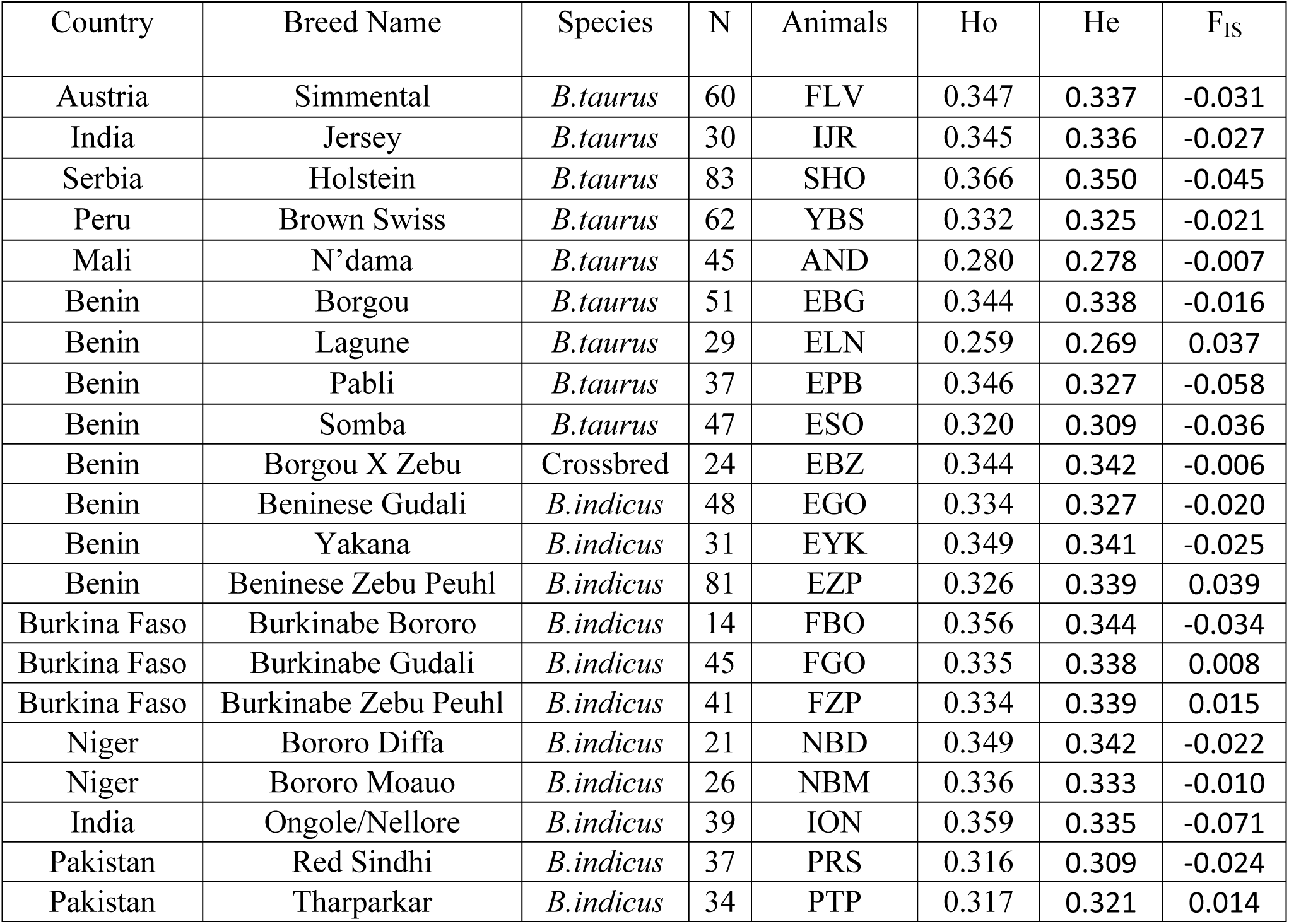
Genetic diversity indicators (Ho, He and F_IS_) for the studies populations. H_o_ is observed heterozygosity, H_e_ is expected heterozygosity, F_IS_ is Wright’s inbreeding coefficient

When comparing the different diversity indices of Beninese zebus with those of South Asian Zebu, we found the estimates were comparable except ION which had H_o_ of 0.359, slightly higher than that of Beninese zebus (EYK, EGO, and EZP). Regarding the H_e_, the EYK and EZP zebus showed the values of 0.341 and 0.339 respectively. Among the zebu, the lowest values of H_o_ and H_e_ were observed in Tharparkar (PTP) and Red Sindhi (PRS) cattle. The inbreeding rate is higher in the Zebu Peuhl of Benin F_IS_ = 0.039 followed by the PTP zebu, F_IS_ = 0.014. The F_IS_ of the EGO, PRS, EYK and ION zebu were negative.

The comparison of the genetic diversity indices between the taurine cattle of Europe and Benin indicated that the highest values of H_o_ and H_e_ were observed in the SHO taurine with respective values of 0.366 and 0.350. The lowest values of these two indices were observed in the ESO and ELN taurine’s. When examining inbreeding coefficients, Benin’s taurine breeds displayed a wider range of values compared to European breeds. The Lagune (ELN) exhibited the highest level of inbreeding (F_IS_ = 0.037) among the Beninese taurine cattle.

The average F_IS_ of taurine’s and zebu of Benin was respectively −0.018 and −0.002 in this study. The F_IS_ varied from −0.058 to 0.037 in the taurine cattle and from −0.0248 to 0.0394 in the zebu cattle. In Benin, the values of H_o_, H_e_ and F_IS_ were higher in the zebu of Benin than in the taurine. The Yakana zebu (EYK) had higher H_o_ and H_e_ values (Ho = 0.349; He = 0.341) while ESO and ELN had the lowest values of these two indices. At the level of the inbreeding coefficient, EZP had the highest value (F_IS_ = 0.039) while ESO and EPB had the respective low values of −0.036 and −0.058.

Among the South Asian and European cattle used as reference breeds in the study, the PRS (H_e_ = 0.316), had the lowest genetic diversity, while the SHO (H_o_ = 0.366) had the highest. F_IS_ values were negative for these reference breeds (except PTP breed) ranging from −0.071 in ION to 0.014 in PTP. Figure 2 shows the distribution of heterozygosity values among different cattle breeds in this study. Each box represents a breed and the vertical axis shows the heterozygosity levels. Lagune cattle breed (ELN) consistently showed the lowest heterozygosity levels suggesting relatively low genetic diversity within the breed. The FBO zebu showed a wide range of heterozygosity values, indicating a diverse genetic composition. In contrast, the AND, EBG and EPB breeds showed moderate levels of heterozygosity, with some individual outliers.

**Figure 1:**
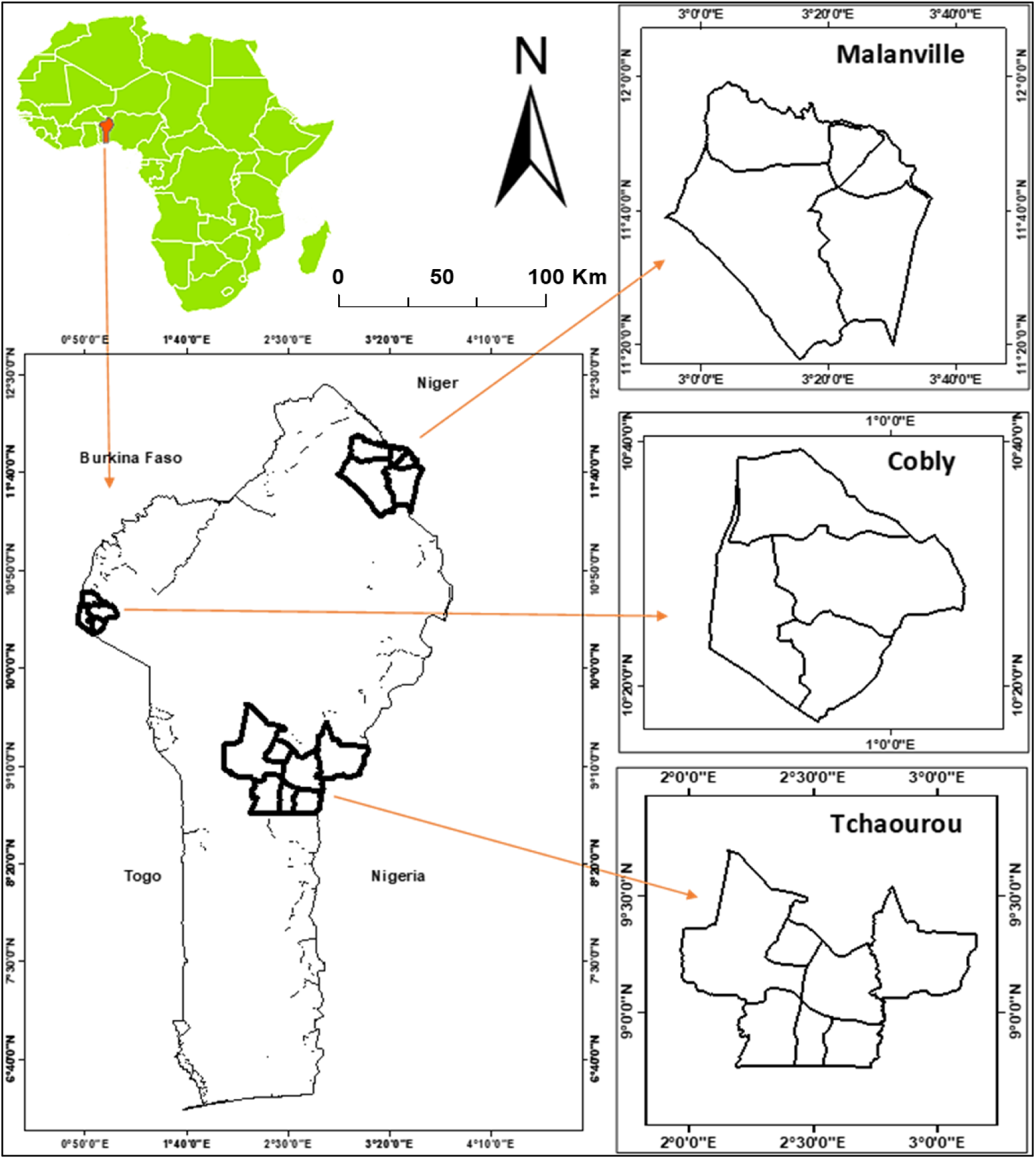
Maps showing the location of Benin cattle breeds under study

**Figure 2:**
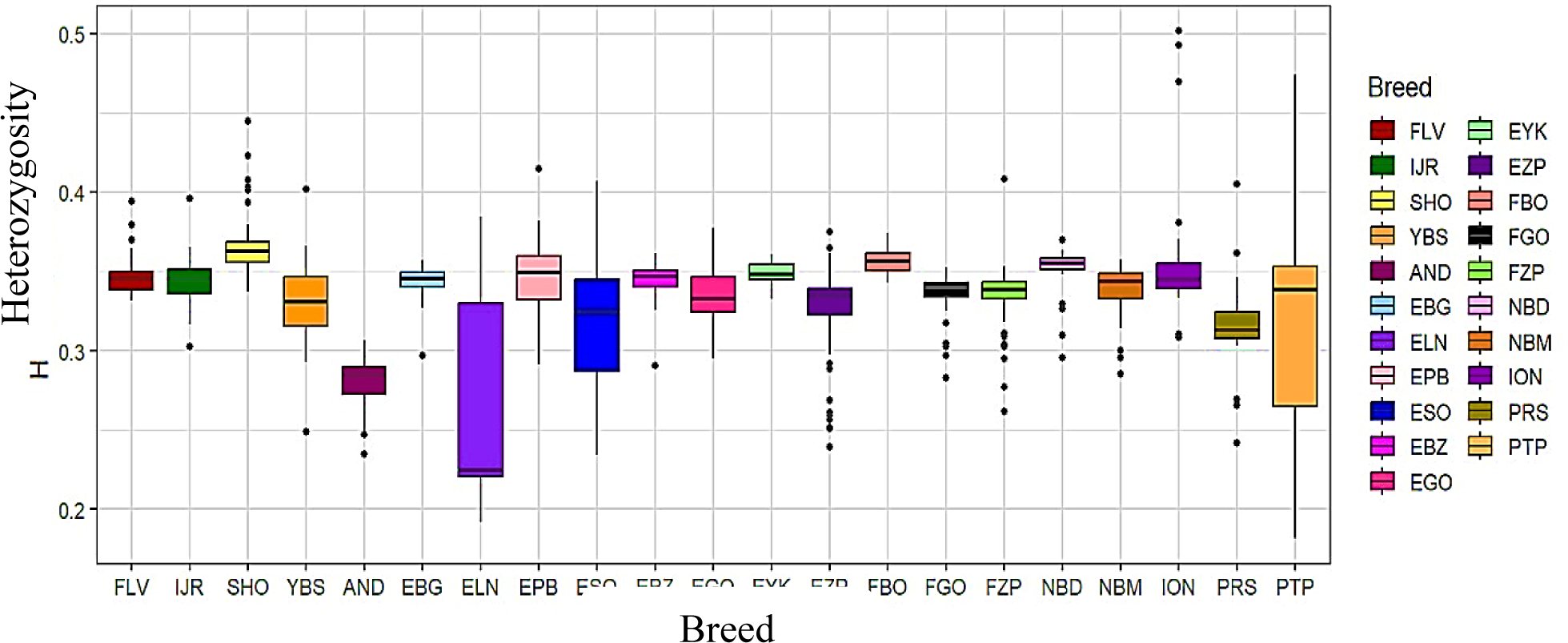
Box plot graph showing the estimates of observed heterozygosity in the breeds under study

### Effective population size (*Ne)*

To estimate the trend in effective population size (*Ne*) for West African taurine and Benin zebu breeds, we used SNeP v1.1, a method based on linkage disequilibrium (LD). The results of the current analysis revealed that the West African taurine breeds exhibited greater genetic variation within their populations compared to the West African zebu breeds (Figure 3). Among the West African taurine breeds (Figure 3A), N’dama (AND) showed a gradual increase in effective population size over the past 750 generations with a peak at around 3000. Borgou (EBG) exhibited a significant *Ne* increase, reaching around 4000. Lagune (ELN) demonstrated an effective population size of around 4000, while Pabli (EPB) showed the highest Ne estimate of approximately 6000. The linkage disequilibrium for these breeds decayed as genetic distance increased, with initial LD values stabilizing around 0.05, indicating recombination over time. For the Beninese zebu breeds, Beninese Gudali (EGO) showed a moderate *Ne* increase over generations, peaking at around 3800 (Figure 3B). Yakana (EYK) exhibited the highest increase, peaking at around 5000. Beninese Zebu Peuhl (EZP) demonstrated a considerable increase, with an effective population size approaching 4000. The LD pattern for these breeds was consistent with that of the taurine breeds, with an initial higher LD that decreased over time, stabilizing around 0.05.

**Figure 3:**
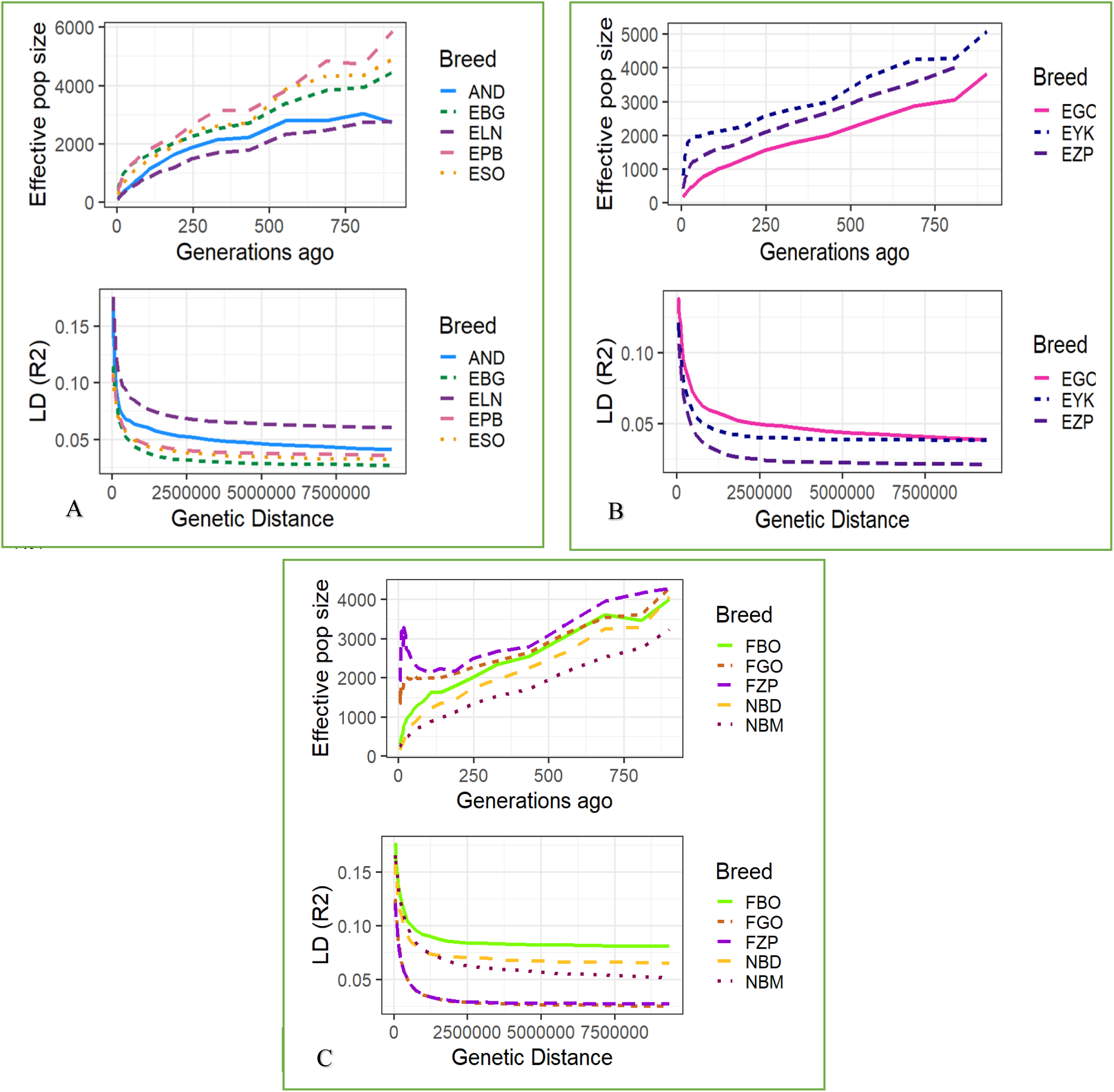
Trends in effective population size of West African taurine and zebu cattle over generations; A-fective population size of West African taurine; B-Effective population size of Benin zebu; C-Effective pulation size of Burkina and Niger zebus.

Among zebu breeds from Burkina Faso and Niger (Figure 3C), the Burkina Bororo (FBO) showed a moderate increase over generations with a peak estimate of about 4000. The Burkinabe Zebu Peuhl (FZP) showed the highest Ne increase with a peaking at over 4000. The Burkinabe Zebu Gudali (FGO) showed a considerable increase, with an effective population size slightly above 4000. NBM showed an effective population size of about 3000, which is the lowest among the studied Zebu populations. Linkage disequilibrium for these breeds (FZP, FGO and NBM) decreased as genetic distance increased, with initial LD values stabilizing around 0.05, indicating recombination over time.

### Genetic relationships and Population structure assessment

Figure 4 displays the results of PCA analysis, which clearly distinguishes European taurines (Simmental, Jersey, Holstein, and Brown Swiss), Asian zebu (Nellore, Red Sindhi, and Tharparkar) and African cattle (N’dama, Pabli, Borgou, Beninese Gudali, Yakana, Zebu Peuhl Benin, Bororo of Burkina Faso, and Niger zebu). Of the overall variation observed in the population, the first principal component (PC1) accounted for 36.75%, while the second principal component (PC2) explained 13.85% of the variation. The PCA results revealed four different clusters of cattle viz. Group1 formed by European taurine cattle, Group2 formed by Asian Zebu, Group3 formed by West African taurine cattle and the Group4 formed by West African Zebu cattle. A range of West African cattle belonging to different populations were located in between Group3 and Group4 clusters indicating varying levels of zebu-taurine admixture in them.

**Figure 4:**
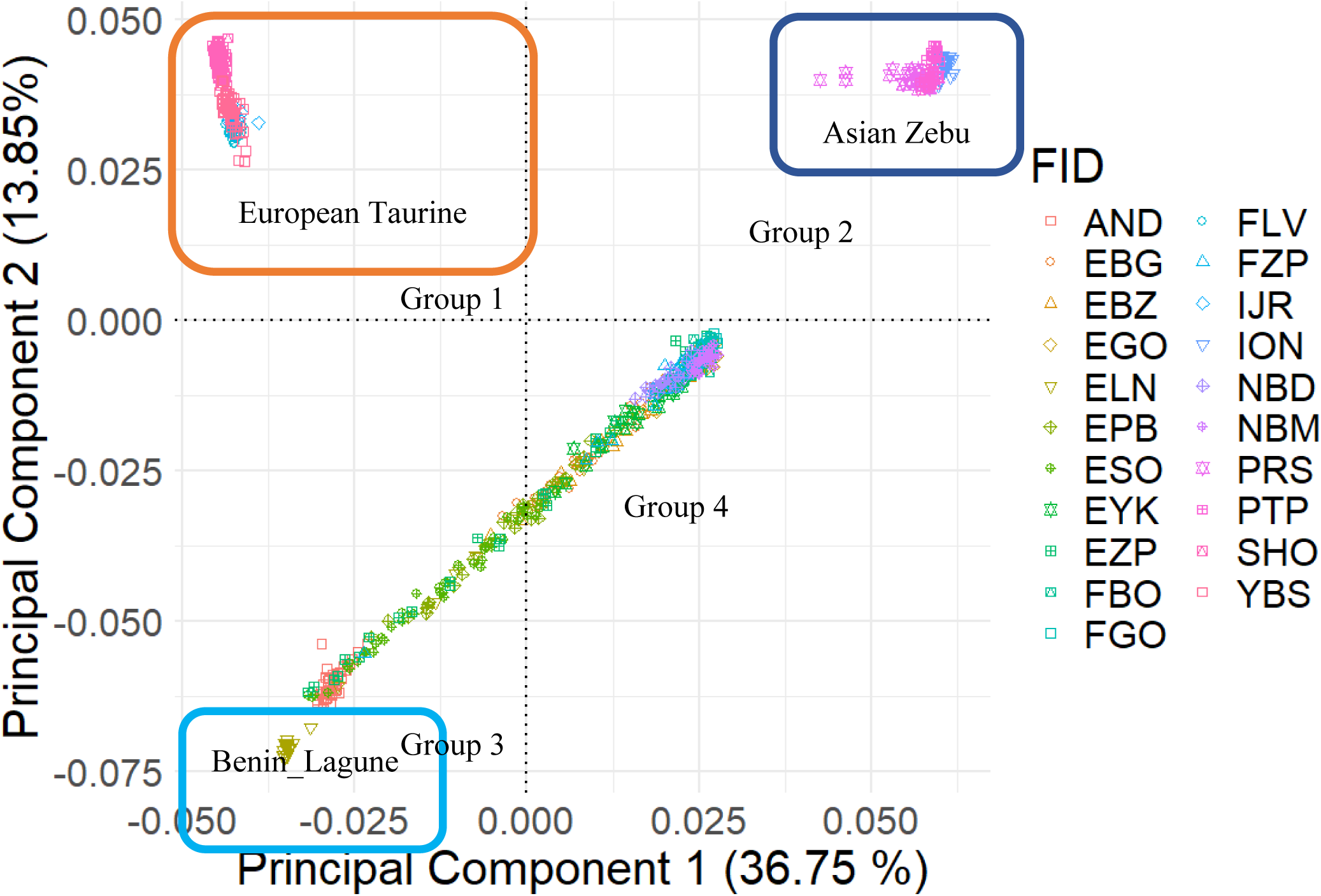
PCA plot obtained from 28.591 SNPs genotypes from 21 cattle populations.

We estimated pairwise Nei’s genetic distance (DG) and visualized the results using a NeighborNet tree (Figure 5). The tree corroborated the results of admixture analysis and showed clear separation of the European taurine and the Asian zebu cattle, highlighting their distinct genetic backgrounds. Within West African cattle populations, NeighborNet analysis revealed several major clusters. The taurine breeds ELN and AND formed a distinct cluster, rooted separately, suggesting greater genetic divergence from the other breeds. Conversely, EYK, EBZ, and EBG appear closely related due to their proximity in the network. Similarly, EGO, FGO, FZP, and EZP formed another closely related cluster, indicating shared ancestral lineages. Interestingly, NBM and NBD showed a notable genetic distance, while FBO and NBM showed closer relationships. The Somba (ESO) and Pabli (EPB) taurine breeds were positioned distantly from other West African taurine’s, suggesting a unique genetic profile arising out of low to moderate zebu admixture in these two breeds.

**Figure 5:**
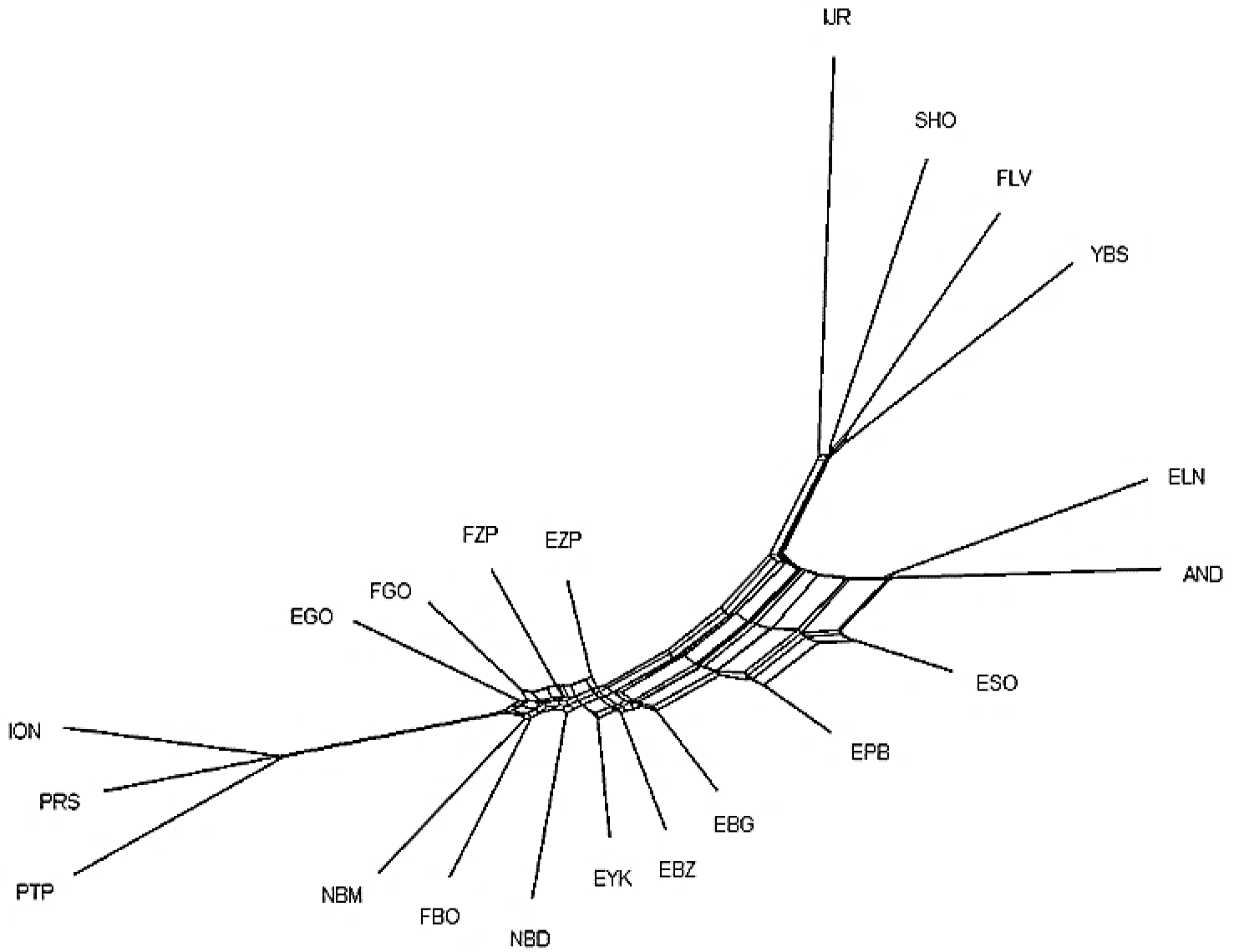
NeighborNet graph estimated from p-distances with SplitsTree

### Admixture and gene flow migration

To investigate the ancestry and genetic admixture in Beninese cattle, an ADMIXTURE analysis was conducted with the number of clusters (K) ranging from 2 to 10 (Figure 6B). The unsupervised clustering revealed a hierarchical ancestry, with the lowest cross-validation error observed at K=10, indicating ten genetic clusters as the most appropriate representation of the data (Figure 6A). Patterns observed at K=2 through K=9 reflected a progressive resolution of breed relationships and the underlying admixture dynamics in West African cattle populations. At K=2, two broad ancestral lineages were evident, distinguishing taurine (*Bos taurus taurus*) from indicine (*Bos taurus indicus*) ancestries. The European taurine breeds (FLV, IJR, SHO, YBS) and South Asian zebu breeds (ION, PTP, PRS) each formed relatively homogeneous clusters, whereas West African breeds displayed heterogeneous ancestry patterns. Among the West African taurines, N’dama (AND) and Lagune (ELN) exhibited predominantly taurine ancestry with little zebu admixture, while Borgou (EBG), Somba (ESO), and Pabli (EPB) showed higher levels of zebu introgression (Figure 4, K=2). At K=3, a distinct African taurine cluster was resolved, separating N’dama (AND) and Beninese taurines (EBG, ELN, ESO, EPB) from European taurines, African zebu and South Asian indicines. Within the Beninese taurine group, Lagune (ELN) showed the lowest proportion of zebu ancestry, Borgou (EBG) the highest, while Somba (ESO) and Pabli (EPB) displayed intermediate levels of admixture.

**Figure 6:**
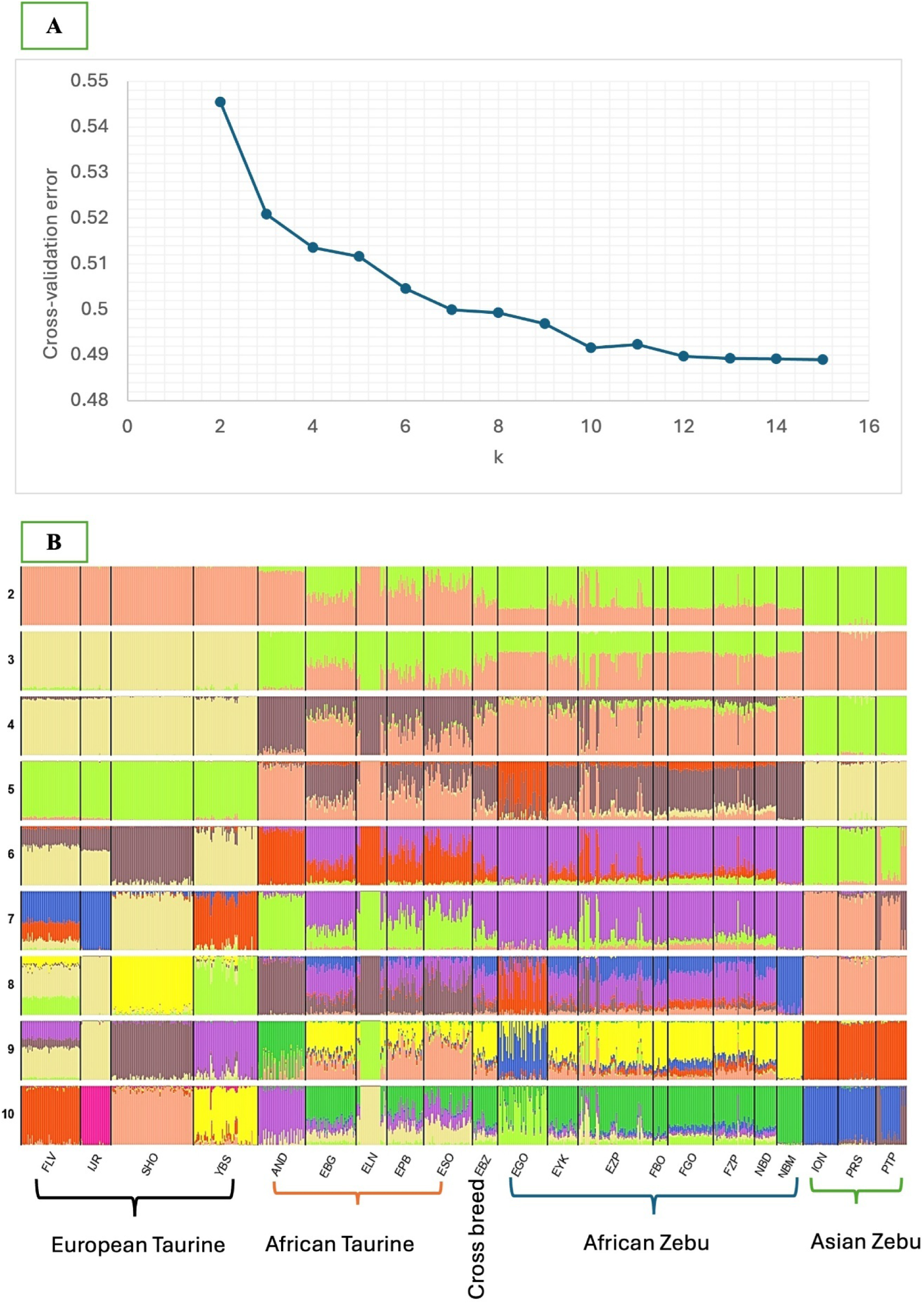
Cross Validation Error plot (**A**). Unsupervised hierarchical clustering of the 852 individuals (**B**). Results for the number of clusters K = 2 to 10 are shown. Individuals are grouped by breed/population. Each individual is represented by a vertical bar. The proportion of the bar in each of K colors corresponds to the average posterior likelihood that the individual is assigned to the cluster indicated by that color. Populations are separated by black lines. Breed abbreviations are as listed in Table 1.

From K=5 onwards, further regional structuring was observed among West African zebu cattle. Beninese Gudali formed a separate cluster at K=5, while Borroro Maoua (NBM) differentiated into an independent cluster at K=8. Among taurine populations, Beninese Lagune (ELN) diverged from N’dama (AND) at K=9, reflecting subtle sub-structuring despite both being predominantly taurine. At the optimal resolution of K=10, N’dama (AND) and Lagune (ELN) formed largely homogeneous clusters with minimal evidence of admixture, whereas Borgou (EBG), Somba (ESO), and Pabli (EPB) showed varying degrees of zebu ancestry. Among West African zebus, Beninese Gudali (EGO) and Nigerien Bororo Maoua (NBM) each formed distinct clusters, while other zebu populations exhibited broadly similar ancestry profiles. Notably, Beninese and Burkinabe Gudali displayed distinct ancestry profiles, whereas Zebu Peuhl from Burkina Faso and Benin shared similar genetic patterns.

The evolutionary relationships and historical gene flow among South Asian Zebu and West African cattle populations were examined using TreeMix, with N’dama (AND) designated as the root (Figure 7). The maximum likelihood tree without migration edges (Figure 7A) depicted the expected divergence between taurine (*Bos taurus taurus*) and indicine (*Bos taurus indicus*) lineages. West African taurines (AND, ESO, EPB and EBG) grouped separately from both West African and South Asian indicines. West African zebu cattle (FGO, FZP, FBO, NBM, EGO, NBD) clustered with South Asian indicines (ION, PRS, PTP), reflecting their shared indicine ancestry (Figure 7A). With incorporation of one migration edge (Figure 7B), gene flow was detected from Pakistani Red Sindhi (PRS) to Pakistani Tharparkar (PTP) cattle. With incorporation of two migration edges (Figure 7C), two additional gene flow events were inferred, both linking Asian indicine lineages to West African zebu: Ongole (ION) to Burkinabe Gudali (FGO) and from Pakistani Tharparkar (PTP) to Yakana (EYK). At three migration edges (Figure 7D), further gene flow was observed from Ongole (ION) to both Red Sindhi (PRS) and Zebu Peuhl (EZP) of Benin. Concurrently, taurine-to-taurine gene flow from N’dama (AND) to Borgou (EBG) was also observed. With four migration edges (Figure 7E), additional indicine introgression was evident. Gene flow from Ongole (ION) extended to Beninese Zebu Peuhl (EZP) and Nigerien Bororo Diffa (NBD), while taurine gene flow from N’dama (AND) to Borgou (EBG) was reaffirmed, indicating sustained influence of N’dama on admixed Beninese taurine’s. At five migration edges (Figure 7F), more complex networks emerged. Borgou (EBG) contributed to Lagune (ELN), while the ancestral junction of N’dama and Lagune was inferred as a source to Pabli (EPB) and Somba (ESO). The same junction also contributed to multiple West African zebu cattle (FBO, NBM, EGO, FGO), suggesting the retention of significant taurine ancestry in them. Further the gene flow observed between Ongole (ION) to Bororo Diffa (NBD), reinforces the repeated input of Asian indicine ancestry across Sahelian zebu populations. The TreeMix analysis revealed the demographic history of West African cattle being shaped by deep taurine–indicine admixture with repeated introductions of Asian indicine ancestry into West African zebu populations. These findings illustrate the impact of long-distance cattle movements, pastoralist exchanges, and crossbreeding practices on cattle genomic diversity in the region.

**Figure 7:**
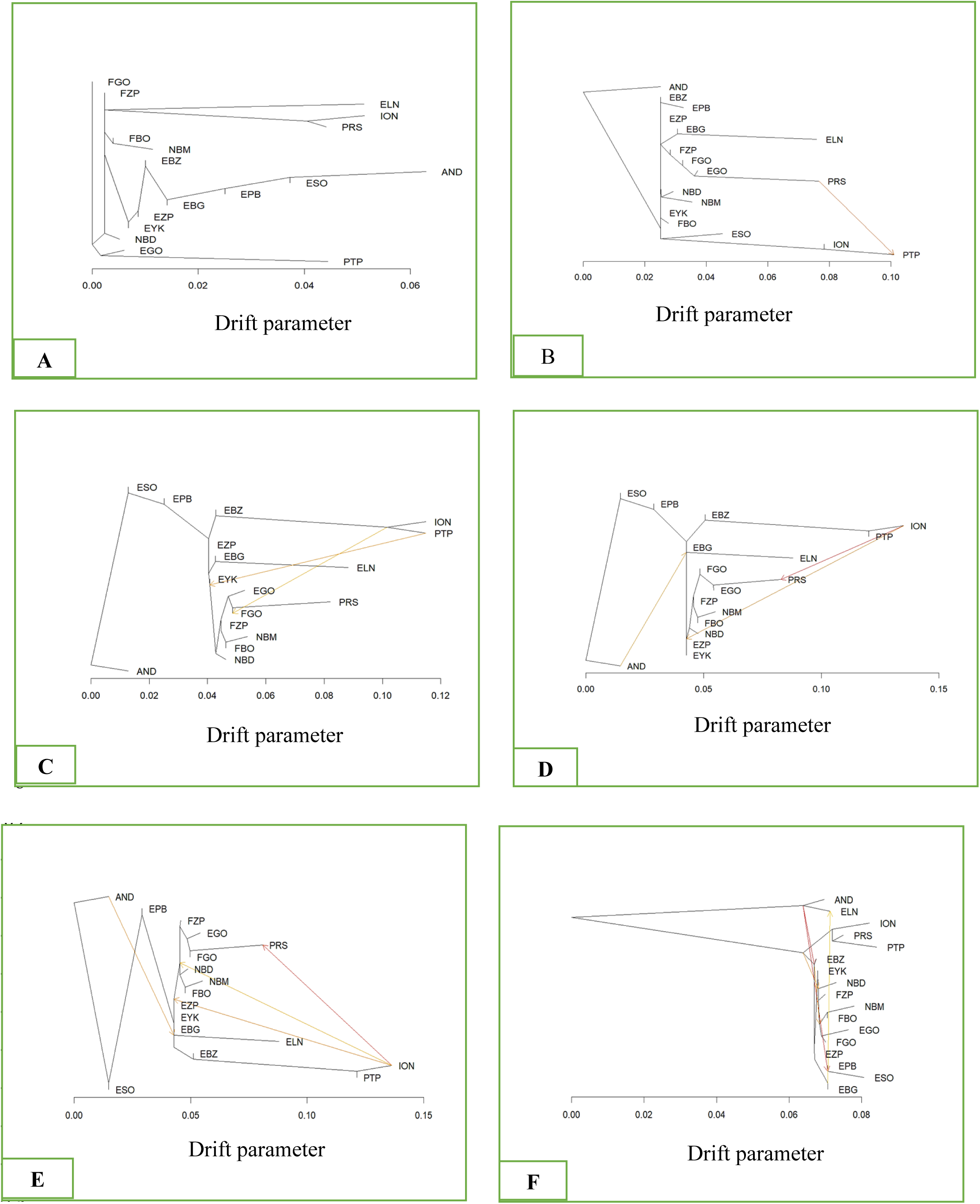
TreeMix analyses depicting evidence of gene flow between different breeds with five asssible migrations by considering N’dama (AND) cattle as root. (A): Maximum likelihood tree. (B-C: TreeMix population splits with one to five migration edges.

The pairwise genetic differentiation statistics (F_ST_) between different cattle breeds investigated in the present study are shown in Figure 8. The genetic differentiation between highly homogenous West African taurine breeds (AND and ELN) and indicine cattle was relatively high, with most pairwise F_ST_ values varying between 0.13 and 0.21. However, Beninese taurine breeds such as Somba, Pabli and Borgou showed low to moderate genetic differentiation with zebu cattle. Among the West African taurine populations, differentiation levels were moderate, with pairwise F_ST_ values ranging between 0.05 and 0.11. N’dama (AND) was consistently the most distinct breed, displaying pairwise F_ST_ values ranging from 0.05 to 0.11 when compared with Beninese taurine populations. This elevated differentiation likely reflects N’dama’s relative geographic isolation and its limited exposure to indicine introgression compared to Beninese taurine cattle. Among the Beninese taurine breeds, Lagune was relatively distinct from Somba (ESO), Pabli (EPB) and Borgou (EBG) with pairwise F_ST_ ranging from 0.05 to 0.11. However, low genetic differentiation was observed between Pabli and Borgou (F_ST_=0.02), Pabli and Somba (F_ST_=0.01) and Borgou and Somba (F_ST_=0.04) suggesting considerable gene flow facilitated by overlapping grazing territories and management practices. Interestingly, the Beninese indicine breed Yakana (EYK) showed no genetic differentiation (F_ST_=0) with Borgou (EBG), Borgou X Zebu crossbred (EBZ) and Beninese Zebu Peuhl (EZP). The differentiation between West African zebu and South Asian indicine cattle was relatively higher with pairwise F_ST_ values ranging from 0.11 to 0.15. Among the West African zebu breeds, differentiation was comparatively low, with pairwise F_ST_ values typically ranging from 0 to 0.03 across Beninese (EGO, EYK, EZP), Burkinabe (FBO, FGO, FZP), and Nigerien (NBD, NBM) populations. These relatively low values indicate that despite their classification as distinct breeds, these zebu populations retain close genetic relationships, likely maintained through transhumance, trade networks, and shared ancestry with South Asian indicines.

**Figure 8:**
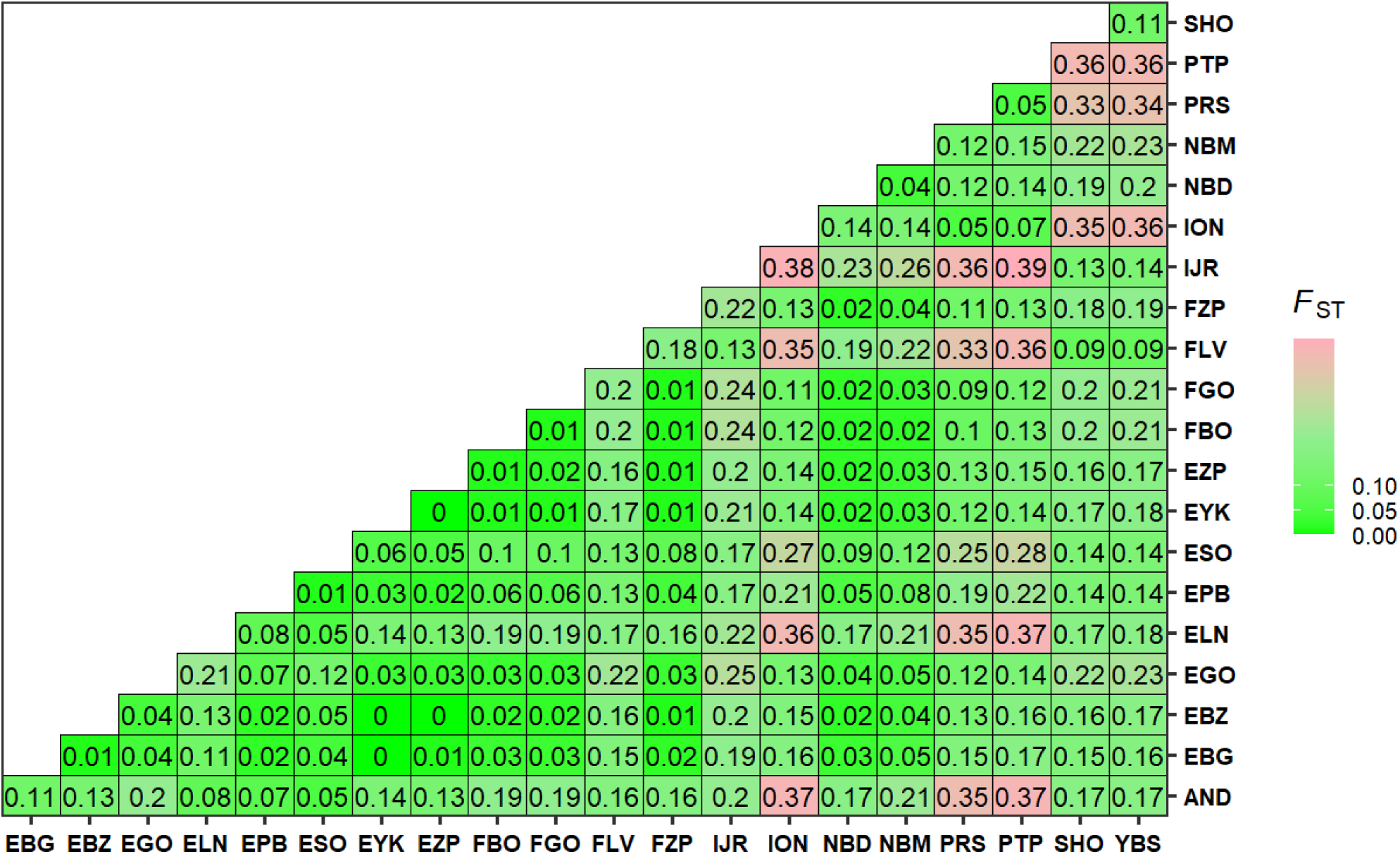
Estimates of the pairwise genetic differentiation statistics (F_ST_)

## Discussion

The analysis of genetic diversity within Benin’s cattle populations provides valuable insights into their population structure, admixture, gene flow, and effective population size (Ne). The combination of these results not only shed light on the historical foundation and genetic variability of local breeds but also reflects the influence of environmental and anthropogenic factors on their genetic makeup. The present investigation into the genetic diversity of native cattle populations of Benin revealed marked differences between taurine and zebu breeds as evidenced by estimated heterozygosity indices. Observed (H_o_) and expected (H_e_) heterozygosity were consistently lower in taurine populations, particularly in the Lagune (ELN) breed, relative to zebu populations such as the Zebu Peuhl. These results are consistent with those reported by [40], who documented H_o_ values ranging from 0.79 in Girolando cattle to 0.57 in the Somba population. In the same study, the highest H_e_ (0.79) was also observed in Girolando cattle, while the lowest (0.63) was recorded in taurine breeds such as Lagune and Somba. The present estimates of heterozygosity are broadly consistent with earlier findings by Moazami-Goudarzi et al. [41] in Somba, Lagune, White Fulani, and Borgou cattle. Álvarez et al. [42] similarly reported comparable heterozygosity levels in various taurine and zebu populations of Burkina Faso. Collectively, these data confirms that West African taurine cattle exhibit relatively low genetic diversity, potentially attributable to long-term geographic isolation, restricted gene flow, and relatively small effective population sizes. In the present study, the observed heterozygosity estimate (H_o_) for Beninese Gudali (0.33) was consistent with earlier report on Cameroonian Gudali (Ho = 0.34)[43] and falling within the same range observed for the Zebu Peuhl (EZP), Borgou (EBG), and Gudali (FGO) cattle. Similar levels of observed heterozygosity, but with slightly elevated estimates were observed in the Pabli (EPB) and Yakana (EYK) populations (H_o_ = 0.35), consistent with reported estimates on Indian (Dixit et al. 2021) and Braizilian breeds of Zebu cattle [44]. Nawaz et al. [45] observed marginally lower H_o_ values in Angus and Hanwoo breeds (0.30 and 0.31, respectively), while similar levels of genetic diversity have also been documented in native Ethiopian cattle populations[46]. In contrast, local Iraqi cattle breeds exhibited higher genetic diversity, with H_e_ values reaching 0.37 [47], exceeding those reported for West African cattle breeds in the present study. Although the H_e_ values observed in this study are lower than those previously reported for the Gudali and Simgud populations of Cameroon[43], they remain substantially higher than those reported for several South African breeds, whose H_e_ ranges between 0.24 and 0.30 [48]. These results highlight an appreciable level of genetic diversity in Gudali cattle from Benin, comparable to that observed in other local African and Asian populations. Such diversity signifies the adaptive potential of these breeds to local environmental conditions and highlights their strategic importance in genetic conservation and sustainable cattle production in sub-Saharan Africa.

The intra-population fixation coefficient (F_IS_) showed considerable variability between the populations studied. Relatively high values were observed in Zebu Peuhl (EZP) and Lagune (ELN) in this study, reaching 0.04, suggesting moderate inbreeding in these groups. Conversely, the negative F_IS_ values recorded in certain taurine and zebu breeds reflect an excess of heterozygotes. This phenomenon could be the result of open community mating practices, commonly found in extensive breeding systems, which tend to limit genetic drift and inbreeding pressure. However, it should be noted that F_IS_ is not a direct measure of the individual inbreeding coefficient. It only reflects the latter in the infrequent case where deviations from Hardy-Weinberg equilibrium are exclusively due to inbreeding [49], [50]. A cautious interpretation of the F_IS_ is therefore essential, taking into account the reproductive dynamics, gene flows and demographic structures specific to each population.

The results from principal component analysis (PCA) confirm the historical genetic divergence between European taurine, Asian zebu, and West African cattle, particularly within Benin’s indigenous breeds. The first two principal components (PC1, 36.75% and PC2, 13.85%) together accounted for more than 50% of the observed genetic variation, highlighting clear clustering of different cattle populations. This genetic structuring is reflective of historical admixture, gene flow events and the selective pressures exerted on these breeds across generations. The Lagune cattle exhibited a distinct genetic identity, showing partial separation from other West African breeds, including those from the neighboring regions. This unique positioning suggests a specific set of adaptive traits shaped by the environmental pressures and breeding practices unique to Benin’s coastal and lagoon regions[24], [51]. The Lagune breed, primarily found in Southern Benin near lagoons and coastal areas, is characterized by its small size, adapted to the constrained environments of its belt[52]. Previous studies have shown that Lagune cattle are genetically distinct from other shorthorn breeds in Africa[41], [53]. Furthermore, Berthier et al.[54] reported a higher degree of trypanotolerance and better resistance to anemia in the Lagune breed compared to the Baoulé breed, further emphasizing its adaptation to local environmental conditions. Historically, the Lagune breed was exported from Benin to various countries in Africa and Europe during the early 20th century, where it was known as the Dahomey cattle[52], [55]. These cattle, maintained by a breeders association in Europe[56], retain many of the phenotypic traits of the original Benin Lagune population, suggesting that the breed has remained genetically stable in Europe. Vanvanhossou et al.[29] further confirmed the purity of this breed by showing that the European Dahomey cattle exhibited over 40% autozygosity, pointing to their shared ancestry with the Beninese Lagune cattle. This finding emphasizes the breed’s genetic integrity despite its long geographical separation.

The results of the principal component analysis also revealed that the majority of West African cattle form a homogeneous group, within which certain individuals belonging to Gudali breed appear dispersed. This distribution suggests that animals phenotypically identified as pure Gudali, pure Borgou,pure Yakana, etc, are probably the result of multiple crosses between the different cattle breeds considered in this study. The results of the principal component analysis (PCA) and admixture analysis corroborate the observations of Opoola et al.[57], who showed that the cattle population of Rwanda exhibits a high degree of crossbreeding, indicating a largely mixed genetic structure. Although misidentification cannot be ruled out, similar observations have been reported in previous work involving East African cattle breeds[58] and native Indian cattle breeds[59], highlighting the limitations of using phenotypic criteria alone to determine breed purity. Furthermore, the ability of PCA to discriminate between groups with marked genetic divergences has been demonstrated in several previous studies[60], [61], [62], [63], [64]. The admixture analysis revealed the extent of gene flow between taurine and zebu populations in West Africa, highlighting the impact of historical and ongoing interbreeding events. At K=2, the analysis uncovers the fundamental genetic separation between *Bos taurus* (European and African taurine) and *Bos indicus* (zebu) cattle. As K values increase, the admixture patterns become more complex. The Lagune cattle (ELN) showed a relatively uniform taurine profile, while other breeds in Benin, including crossbred zebu populations showed significant taurine-zebu admixture. This admixture is consistent with the observation that zebu cattle, especially from Asia, were introduced in West Africa centuries ago and interbred with local taurine populations [65]. Furthermore, admixture patterns in EBG, EPB, and ESO breeds suggest that breeding strategies, possibly influenced by the movement of Fulani herders and their zebu cattle, have reinforced zebu traits that confer advantages in the region’s challenging climate, including heat tolerance and disease resistance[66], [67]. The presence of zebu introgression in some West African cattle populations is also supported by the study on the geographic spread of zebu cattle, which were originally domesticated in the Indus Valley and later migrated to Africa[31]. The work of Koudandé et al.,[68] revealed, via Y chromosome analysis, a marked zebu introgression, ranging from 37.5% in the Benin Valley and Plateau East to 100% in the Plateau West. Admixture analysis (k = 2) confirms the widespread presence of zebu genes in the populations studied[68]. The results of Kassa Sagui et al.[40] also confirmed an average zebuine introgression of 20% within local taurine breeds, highlighting the urgent need to set up a structured genetic improvement program to preserve Benin’s indigenous taurine cattle heritage.

Gene flow migration analysis, using TreeMix, further elucidated the dispersal of Zebu genetics across native cattle populations. Notably, migration events from Asian zebu populations, such as ION to the Gudali zebu contributed to the genetic makeup of Benin cattle. These results underscore the significant role of external gene flow in shaping the genetic diversity of Benin’s indigenous cattle breeds. Migration events from taurine breeds like AND and ELN into Borgou (EBG) further reinforce the genetic contribution of local African taurine populations in the region. As migration events increase, additional flows from taurine N’dama (AND) to the Lagune breed (ELN) and between taurine populations (AND to EPB and ESO) highlighted the interconnectedness of Benin’s cattle breeds. These gene flow patterns not only point to a history of genetic exchange but also suggest ongoing hybridization, with taurine traits being incorporated into the local zebu populations [69]. These migration events play a crucial role in enriching the genetic pool, conferring resilience to environmental stresses such as disease, heat, and feed scarcity.

The effective population size (N_e_) provides a critical measure of the genetic health and diversity of cattle populations. The relatively high N_e_ values observed in taurine breeds such as Pabli and Lagune (∼6000 and ∼4000, respectively) indicate substantial genetic variability, likely due to less restrictive mating systems and larger population sizes in more remote areas [26]. These high N_e_ values suggest that these breeds are not only genetically diverse but also resilient to the pressures of inbreeding. The N_e_ values for Benin’s zebu breeds are moderately high, with peaks around 5000 in the Yakana zebu, indicating a more controlled but substantial genetic diversity. This controlled diversity helps to maintain a stable genetic base that can adapt to selective pressures while preventing excessive inbreeding and loss of genetic diversity.

The genetic differentiation and admixture patterns observed in the Benin cattle populations provide compelling insights into the evolutionary history and genetic relationships between the indigenous taurine and zebu breeds. The Pabli breed, which was historically reported in the Kerou region of Northwest Benin, showed slight genetic differences from the Borgou breed despite being considered extinct due to crossbreeding[70], [71]. Recent findings, however, have revealed that small populations of Pabli cattle still exist, exhibiting distinct genetic characteristics that can be preserved as a valuable genetic reservoir for future breeding efforts. The pairwise F_ST_ values, highlight the varying degrees of genetic divergence across the different populations. For instance, the lack of differentiation (F_ST_ = 0) between the EBG, EYK, and EZP populations suggests substantial gene flow or a shared evolutionary history, likely driven by historical interactions or crossbreeding practices. In contrast, the moderate F_ST_ values of 0.11 observed between Borgou and N’Dama breeds, suggest selective pressures, local environmental adaptation, or geographical isolation contributing to genetic differentiation, which is consistent with the differentiation of taurine breeds seen across West Africa[72]. This trend is further emphasized by the NeighborNet analysis, where taurine breeds such as ELN and AND clustered separately from European taurine breeds, confirming the role of historical and geographical factors in shaping the genetic profiles of these populations[58], [73]. Furthermore, the moderate differentiation observed between Benin’s zebu populations and those from neighboring countries such as Niger and Burkina Faso (F_ST_ = 0.01–0.05) indicates regional gene flow, likely facilitated by transhumance pastoralism, which is a common practice in West Africa. However, the higher differentiation between EGO (Benin) and NBM (Niger) populations (F_ST_ = 0.05) suggests the influence of localized breeding systems and environmental factors on genetic divergence.

The admixture analysis further supports these findings, with the Lagune (ELN) breed displaying a taurine genetic profile, while other Benin zebu populations show significant taurine-zebu admixture, reflecting selective pressures such as heat tolerance and disease resistance in response to the challenging West African Sahel environment[73], [74]. This genetic complexity is mirrored by studies in other parts of Africa, such as those involving zebu and taurine breeds in East and Central Africa, where F_ST_ values below 0.01 suggest frequent gene flow and low differentiation within subpopulations[46], [75]. The moderate differentiation observed between East and West African cattle populations, such as the N’Dama and Muturu breeds, further supports the hypothesis that geographical and environmental factors influence genetic divergence in a manner similar to what is seen in Benin’s indigenous cattle[75]. Moreover, the westward movement of zebu cattle across Africa, potentially along historical trade routes, would have contributed to the decreasing zebu ancestry observed from East to West Africa, as described by Hanotte and Kambal[76]. These findings emphasize the dynamic and ongoing nature of cattle population genetics in Africa, with the complex interactions between indigenous taurine cattle and zebu playing a central role in shaping the genetic diversity observed in Benin’s cattle populations. The study’s results thus underline the necessity for region-specific breeding management strategies to preserve genetic diversity, particularly in the face of environmental and anthropogenic pressures, ensuring the sustainability and resilience of these cattle populations.

## Conclusion

This study provides an in-depth analysis of genetic diversity, population structure, and gene flow within indigenous cattle breeds of Benin. The results highlighted significant genetic diversity, particularly in taurine breeds such as Lagune and Pabli, reflecting their resilience and adaptive potential. Admixture analysis revealed significant zebu introgression in several taurine populations, reflecting historical and recent crossbreeding with Asian and African zebus, particularly facilitated by transhumance. These results underscore the importance of taking genetic variability and introgression dynamics into account in any genetic management or improvement strategy. On a practical level, the knowledge gained from this study can serve as a basis for the development of Community Based Breeding Programs (CBBPs) adapted to local contexts, integrating breeders’ preferences, productive performance, and the conservation of genetic diversity. Such programs are essential to ensure the sustainability of cattle production systems in West Africa by promoting controlled crossbreeding and genetic management plans based on robust scientific data. Future research could The present study constituted a key step toward the sustainable valorization of local bovine genetic resources in response to the growing challenges of productivity, resilience, and food security in West Africa.

## Acknowledgments

The authors express their gratitude to all breeders for their valuable collaboration throughout the study. We also extend our sincere thanks to the entire staff of the Animal Production and Health Laboratory (APHL) in Seibersdorf, Vienna, for their unwavering support, dedication, and kindness.

## Author contributions

LZ, KP, ITA and IH designed the study. LZ and RP performed laboratory analysis. LZ, TKM, SRT and NT conducted data analysis. LZ wrote the first draft of the manuscript. KP, IH, ASA, HSSW, MA, CI and AT critically reviewed the manuscript. KP, ITA and IH supervised the study. All authors read and approved the final manuscript.

## Additional Information

Supplementary Information S1 (BEN_LOUK_BIODIV_FINAL_GENOTYPES.txt)

